# A single-cell reference atlas delineates CD4^+^ T cell subtype-specific adaptation during acute and chronic viral infections

**DOI:** 10.1101/2021.09.20.458613

**Authors:** Massimo Andreatta, Zachary Sherman, Ariel Tjitropranoto, Michael C. Kelly, Thomas Ciucci, Santiago J. Carmona

## Abstract

CD4^+^ T cells are critical orchestrators of immune responses against a large variety of pathogens, including viruses. The multifaceted roles of CD4^+^ T cells, including their help to innate cells, CD8^+^ T and B cells and their support for long-lived immunity rely on a profound functional heterogeneity. While multiple CD4^+^ T cell subtypes and their key transcriptional regulators have been identified, there is a lack of consistent definition for CD4^+^ T cell transcriptional states. In addition, the progressive changes affecting CD4^+^ T cell subtypes during and after immune responses remain poorly defined. Taking advantage of single-cell transcriptomics, efficient computational methods, and robust animal models, we characterize the transcriptional landscape of CD4^+^ T cells responding to self-resolving and chronic viral infections. We build a comprehensive atlas of virus-specific CD4^+^ T cells and their evolution over time, and identify six major distinct cell states that are consistently observed in acute and chronic infections. During the course of acute infection, T cell composition progressively changes from effector to memory states, with subtype-specific gene modules and kinetics. Conversely, T cells in persistent infections fail to transition from effector to memory states, and acquire distinct, chronicity-associated transcriptional programs. By single-cell T cell receptor (TCR) sequencing analysis, we characterize the clonal structure of virus-specific CD4^+^ T cells across individuals and T cell subtypes. We find that virus-specific CD4^+^ T cell responses are mainly private across individuals and that most T cells differentiate into all subtypes independently of their TCR, in both acute and chronic infections. Finally, we show that our CD4^+^ T cell atlas can be used as a reference to accurately interpret cell states in external single-cell datasets. Overall, this study describes a previously unappreciated level of adaptation of the transcriptional states of CD4^+^ T cells responding to viruses and provides a new computational resource for CD4^+^ T cell analysis, available online at https://spica.unil.ch.

## Introduction

CD4^+^ T cells play a critical role in shaping immune responses against pathogens through the secretion of soluble mediators and direct cell interactions with other immune cell populations. The multifaceted ability of CD4^+^ T cells to orchestrate multiple layers of protection relies on their unique capacity to adopt diverse functional fates upon antigen encounters (*Nguyen et al., 2019; Swain et al., 2012*). Following viral infections, naive CD4^+^ T cells clonally expand and differentiate into effector populations supporting both cellular and humoral responses. This functional diversification of CD4^+^ T cell populations is under tight transcriptional control, ensuring the appropriate positioning and deployment of effector functions (*Zhu et al., 2010*). While Th1 cells, supported by the transcription factors Blimp1 and Tbet, regulate cellular responses in helping CD8^+^ T cells and innate populations through the secretion of INF-γ, follicular-helper CD4^+^ T cells (Tfh), which depend on Bcl6, promote antibody responses via direct cell contact with B cells and the production of cytokines like IL-21 (*Crotty, 2011; Laidlaw et al., 2016; Sheikh and Groom, 2021*).

During the contraction phase following its initial amplification, the pool of virus-specific CD4^+^ T cells declines in both self-resolving and persistent infections. However, the nature of the infection greatly impacts the evolution of CD4^+^ T cell functions (*Brooks et al., 2005; Crawford et al., 2014; Fahey et al., 2011*). After acute viral infections, pathogen clearance is followed by the persistence of memory CD4^+^ T cell populations that acquire distinct phenotypes, gene expression and functional properties (*Crawford et al., 2014; Hale et al., 2013; Marshall et al., 2011*). Because memory populations are heterogeneous and include many subsets, including Th1- and Tfh-like subsets, measuring transcriptional changes occurring between the effector to memory populations remains challenging. In fact, in addition of Th1 and Tfh cells, memory populations are comprised of a less differentiated subset of Central Memory (Tcm) cells that contribute to long-term protective functions of CD4^+^ T cells (*Pepper and Jenkins, 2011*). Tcm phenotypically resemble cells present during the early anti-viral response and referred to as Central Memory precursors (Tcmp), raising the possibility of an early transcriptional imprinting that favors the emergence of long-lived memory CD4^+^ T cells (*Ciucci et al., 2019; Marshall et al., 2011; Pepper et al., 2011*). Yet, the nature of such program, as well as its overlap with that involved in Th1 and Tfh subsets, has not been elucidated. More broadly, it remains unclear whether a shared transcriptional module regulates memory differentiation, or whether diverse gene programs allowing long-term maintenance are imprinted in a subset-specific manner. In sharp contrast to acute settings, chronic infections do not result in such phenotypic memory transition of persisting cells (*Brooks et al., 2005; Fahey et al., 2011*). Instead, in response to sustained antigenic stimulation, CD4^+^ T cells acquire dysfunctional features, including the expression of inhibitory receptors and reduced cytokine production (*Brooks et al., 2005; Crawford et al., 2014*). In this context, questions remain about how persistent infections alter the functional and transcriptional landscape of CD4^+^ T cell populations. However, because of the lack of consistent definition of virus-specific CD4^+^T cell states across conditions and over time, the subtype-specific adaptations during infections are currently poorly characterized.

Although there is evidence that cell fate decisions are stochastically imprinted on T cells (*Buchholz et al., 2013; Buchholz et al., 2016; Soon et al., 2020*), other studies have shown that cell-intrinsic factors as well as environmental cues affect the differentiation of single naïve T cells (*Cho et al., 2017; Tubo et al., 2016; Tubo et al., 2013*). Among these factors, the interaction between the T cell receptor (TCR) and their cognate antigen has been shown to influence the diversification and maintenance of CD4^+^ T cells (*Cho et al., 2017; Snook et al., 2018; Tubo et al., 2013*). For instance, recent studies showed that Th1 and Tfh differentiation are influenced by TCR usage, affinity to cognate peptides and the type of infection (*Khatun et al., 2021; Kunzli et al., 2021; Snook et al., 2018*). Yet, we do not fully understand the extent by which the TCR repertoire impacts the early fate decision of CD4^+^ T cells responding to acute and chronic viral infections. In particular, it remains to be addressed whether naïve T cells with particular TCR chains are preferentially recruited during the effector phase and adopt specific transcriptional profiles that could skew the overall immune response.

Here we employed single-cell RNA sequencing (scRNA-seq) coupled with single-cell TCR sequencing to explore the landscape of virus-specific CD4^+^ T cell states at different timepoints during acute and chronic infections. We provide evidence of both shared and subtype-specific transcriptional changes occurring dynamically in both types of infection. Analysis of paired scRNA-seq and scTCR-seq data of antigen-specific polyclonal T cells revealed that, although a fraction of the clonotypes in both acute and chronic settings were significantly biased towards specific subtypes, T cell functional diversification appears to be mostly independent of the expression of particular TCR chain pairs. Based on these results, we make available a new reference atlas describing virus-specific CD4^+^ T cell states – including Th1, Tfh and Tcmp/Tcm – and their dynamic evolution over time during acute and chronic infection. By combining this atlas with a reference-projection algorithm, we provide a new computational framework that enables automated and accurate interpretation of CD4^+^ T cell states across models, conditions and experiments.

## Results

### Differential phenotypic adaptation of CD4^+^ T cells in acute and chronic viral infection

To characterize the diversification of T cell populations during acute and chronic infections, we used two variants of the lymphocytic choriomeningitis virus (LCMV): the Armstrong and Clone 13 strains. Both viruses induce a strong T cell amplification early in the response, followed by the persistence of a small pool of virus-specific T cells at later timepoints. While the Armstrong strain results in an acute infection cleared within 6-8 days post infection (dpi), Clone 13 persists, leading to chronic infection (*Ahmed et al., 1984; Crawford et al., 2014*). Using these models, we sought to measure the phenotypic changes of virus-specific T cells following infection, both at early and late timepoints. Virus-specific CD4^+^ and CD8^+^ T cells were identified using MHC tetramer loaded with the LCMV-derived GP66 and GP33 peptides, respectively. Cells were analyzed using spectral flow cytometry with a panel of 21 parameters allowing high-dimensional analyses based on 14 surface markers expressed by virus-specific T cells (***Fig. 1A, S1A-C***).

**Figure 1:**
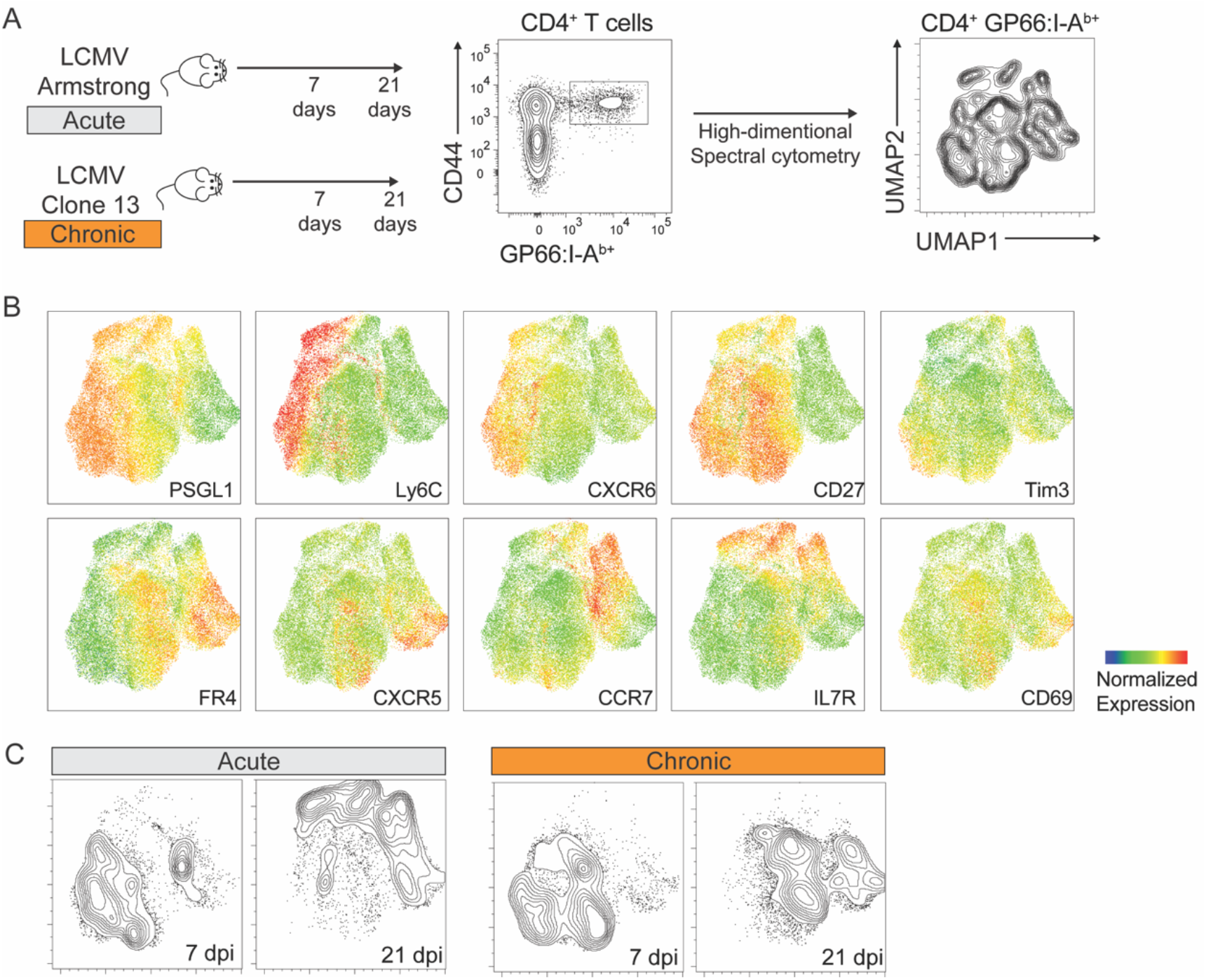
Phenotypic characterization of virus-specific CD4^+^ T cells by spectral flow cytometry. GP66:I-A^b+^ CD4^+^ T cells were analyzed 7 and 21 days after infection with LCMV Armstrong and Clone 13. **A)** Schematic of experimental procedures. Uniform Manifold Approximation and Projection (UMAP) visualization was calculated based on the expression of 14 markers on virus-specific CD4^+^ T cells. **B)** Expression of selected markers shown on the UMAP as in A). **C)** CD4^+^ T cells from each condition were highlighted as contour lines on the UMAP. Experiment with 4-5 mice per group, representative of 2 independent experiments.

Cytometric readouts showed that virus-specific CD4^+^ T cells are highly heterogeneous and largely differ phenotypically from CD8^+^ T cells, both at early and late timepoints after acute and chronic infections (**Fig. 1B**, ***S1D***). Interestingly, while CD4^+^ T cells responding to acute and chronic infections at early timepoints showed partial similarities, minimal overlap was observed at late timepoints. Additionally, CD4^+^ T cells differentiating in chronic settings appear to change less drastically over time compared to the sharp transition occurring between early and late timepoints after acute infection (***Fig. 1C***). Overall, these analyses revealed a profound and fast-adapting phenotypic heterogeneity of CD4^+^ T cell populations in response to different infection settings.

### Defining the landscape of CD4^+^ T cell states in acute and chronic viral infection

To gain further insight into the heterogeneity and transcriptional adaptation of CD4^+^ T cells, virus-specific GP66:I-A^b+^ were purified from LCMV-infected animals either 7 or 21 days after Clone 13 infection, conditions referred to as Early and Late Chronic. In addition, similar populations were isolated 7, 21 and >60 days after LCMV Armstrong infection (samples referred to as Acute, Early and Late Memory, respectively). A total of 11 datasets were processed with droplet-based single-cell RNA sequencing resulting in over 35,000 high-quality virus-specific CD4^+^T cell transcriptomes **(*Fig. 2A***). In addition, selected samples were used to measure TCR usage and transcriptome simultaneously at the single-cell resolution **(*Supplemental Data 1***). To generate a unified atlas of virus-specific CD4^+^ T cell states in acute and chronic infections, all datasets were integrated with STACAS, a computational tool allowing correction of batch effects while preserving relevant biological variability across datasets (*Andreatta and Carmona, 2021a*) (see Methods) (***Fig. 2B***). Importantly, while different timepoints and types of infection (i.e. acute vs chronic) occupied different areas of the integrated space, biological replicates were largely covering overlapping areas of the atlas (***Fig. S2A***), confirming a successful data integration. By clustering the high-dimensional space of the integrated atlas, we defined six major and three minor CD4^+^ T cell clusters that were annotated based on the expression of canonical markers previously described in this model. Among the six major clusters, representing >93% of the cells in the atlas, we identified: *i)* Th1 effector cells, expressing the highest levels of *Cxcr6* and *Ly6c2* (encoding Ly6C); *ii)* Tfh effector cells, preferentially expressing *Cxcr5* and *Izumo1r* (encoding FR4); *iii)* Central memory precursors (Tcmp) expressing the highest levels of *Ccr7* (***Fig. 2C***). The remaining three major clusters were identified as putative memory populations based on their higher expression of memory-associated genes *Tcf7* (encoding TCF1) and *Il7r*, corresponding to *iv)* Th1 memory (co-expressing *Tcf7*, *Il7r*, *Cxcr6* and *Ly6c2*), *v)* Tfh memory (co-expressing *Tcf7*, *Il7r* and *Izumo1r*) and *vi)* Central Memory cells (Tcm), with the highest levels of *Tcf7* and *Il7r* but limited expression of Th1/Tfh marker genes (***Fig. 2C***). Consistent with these annotations, Th1 memory, Tfh memory and Central Memory (Tcm) populations were predominantly derived from virus-specific CD4^+^ T cells isolated at late timepoints after acute infection (***Fig. S2A***, see next section). In addition, these clusters of virus-specific CD4^+^ T cells and their annotations were independently validated using gene signatures we previously identified on CD4^+^ T cells after acute viral infection (*Ciucci et al., 2019*) (***Fig. S2B***). Finally, three minor clusters corresponded to *i) Foxp3-expressing* regulatory T cells (Treg), *ii)* a Tfh-like state expressing high levels of type 1 interferons-stimulated genes (INFI-stimulated) and *iii)* a population characterized by high levels of *Eomes* (Eomes-HI) (***Fig. 2C***). These minor states were largely associated to chronic infection and will be described in a later section.

**Figure 2:**
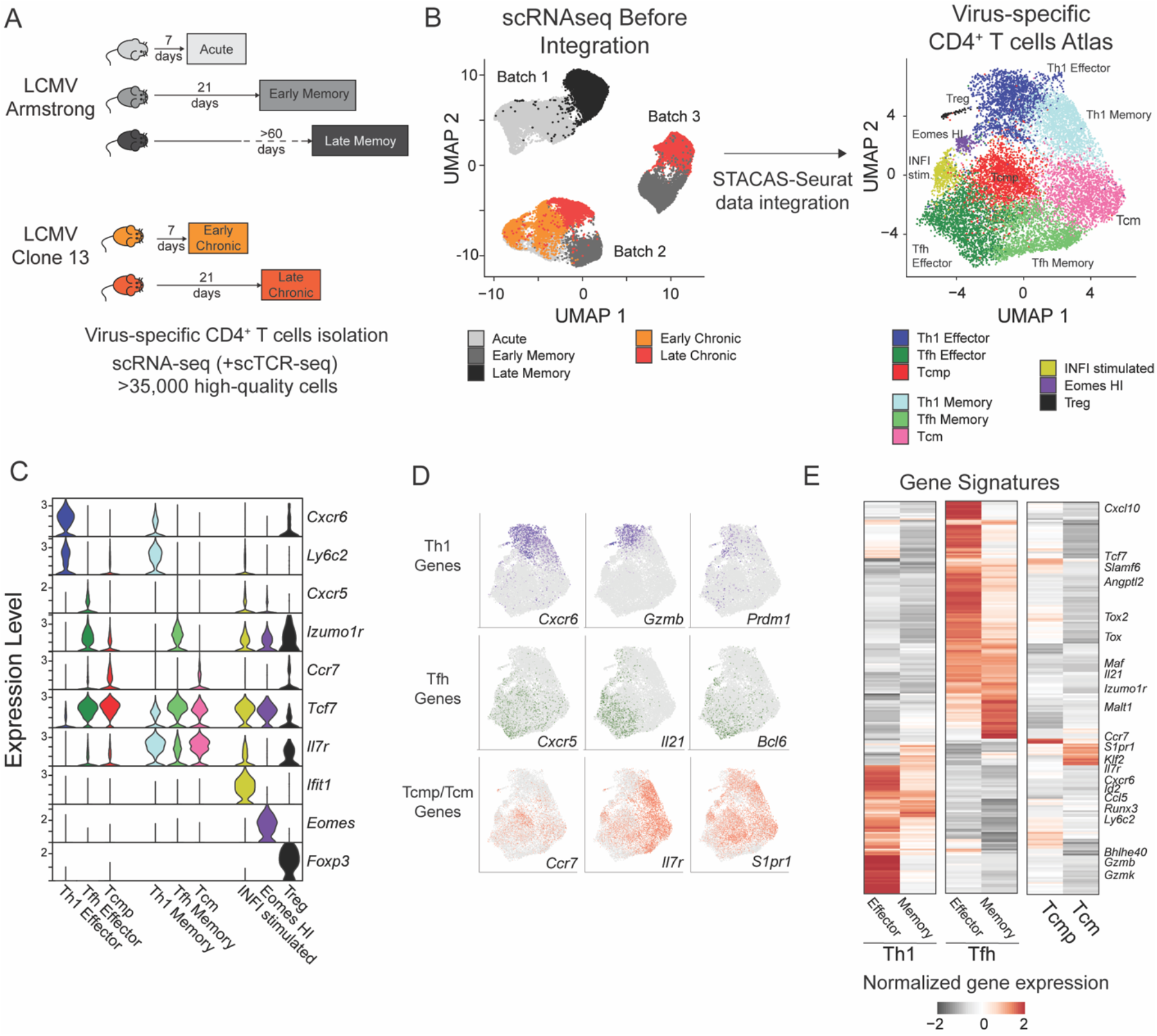
Transcriptional landscape of CD4^+^ T cell states during infections. **A)** Schematic experimental design to assess virus-specific T cell transcriptomes at different timepoints in acute (Armstrong) and chronic (Clone 13) LCMV infections. **B)** UMAP visualization of single-cell data before and after dataset integration, highlighting the samples and batches on the left and the 9 CD4^+^ T cell subtypes of the reference atlas on the right. **C)** Expression levels of key marker genes in the 9 subtypes of the reference atlas. **D)** Single-cell expression visualized in the UMAP space for key marker genes of Th1, Tfh and Tcmp/Tcm subtypes. **E)** Average expression of differentially expressed genes in the six major subtypes of the reference atlas; selected genes are highlighted.

We next explored the expression of genes involved in the function of CD4^+^ T cell subsets across major clusters. As expected, Th1 cells, both effector and memory, expressed the highest amount of the Th1-defining transcription factor *Prdm1* (encoding Blimp1) together with *Gzmb –* encoding the cytotoxic effector molecule Granzyme B. Similarly, the Tfh-specific transcription factor *Bcl6* and effector molecule *Il21* were almost exclusively expressed in Tfh cells (***Fig. 2D***). Interestingly, Tcmp and Tcm populations, which expressed the highest levels of *Ccr7*, *Il7r* and *S1pr1* were characterized by the promiscuous expression of both Th1-associated genes such as *Nkg7*, *Ifngr1* or *Runx3*, and Tfh-specific markers like *Slamf6*, *Tox* or *Bcl2* (***Fig. 2CD, S2C***).

Differential gene expression analysis between the major CD4^+^ T cell states revealed additional subtype-specific genes, and showed that Th1 states (both Effector and Memory subsets) share a common gene module (e.g. *Id2*, *Runx3*, *Gzmb*), distinct to that of Tfh cells, characterized by the expression of *Tox*, *Maf* and *Izumo1r* (***Fig. 2E***). Although Tcmp and Tcm states are largely defined by their lack of Th1 and Tfh gene signatures, consistent with a more quiescent, undifferentiated state, they are characterized by the shared expression of genes such as *Ccr7*, *Klf2* and *S1pr1*. The full list of CD4^+^ T cell subtype-specific signatures is available in ***Supplemental Data 2***, and gene expression in this dataset can be explored online at https://spica.unil.ch/refs/viral-CD4-T.

### Subtype-specific evolution of CD4^+^ T cell states to acute infection

To further describe CD4^+^ T cell states as they adapt during the course of an acute, self-resolving infection that generates protective memory T cells, we investigated subtype composition and subtype-specific transcriptional changes over time. First, from the effector phase (7 dpi) to the early memory phase (21 dpi), we observed a dramatic shift from Th1 effector, Tcmp and Tfh effector to Th1 memory, Tcm and Tfh memory **(*Fig. 3A*)**. This was consistent with the fact that major functional and phenotypic changes occurs during and after the contraction phase 12-20 days following infection (*Marshall et al., 2011; Pepper et al., 2011*).

**Figure 3:**
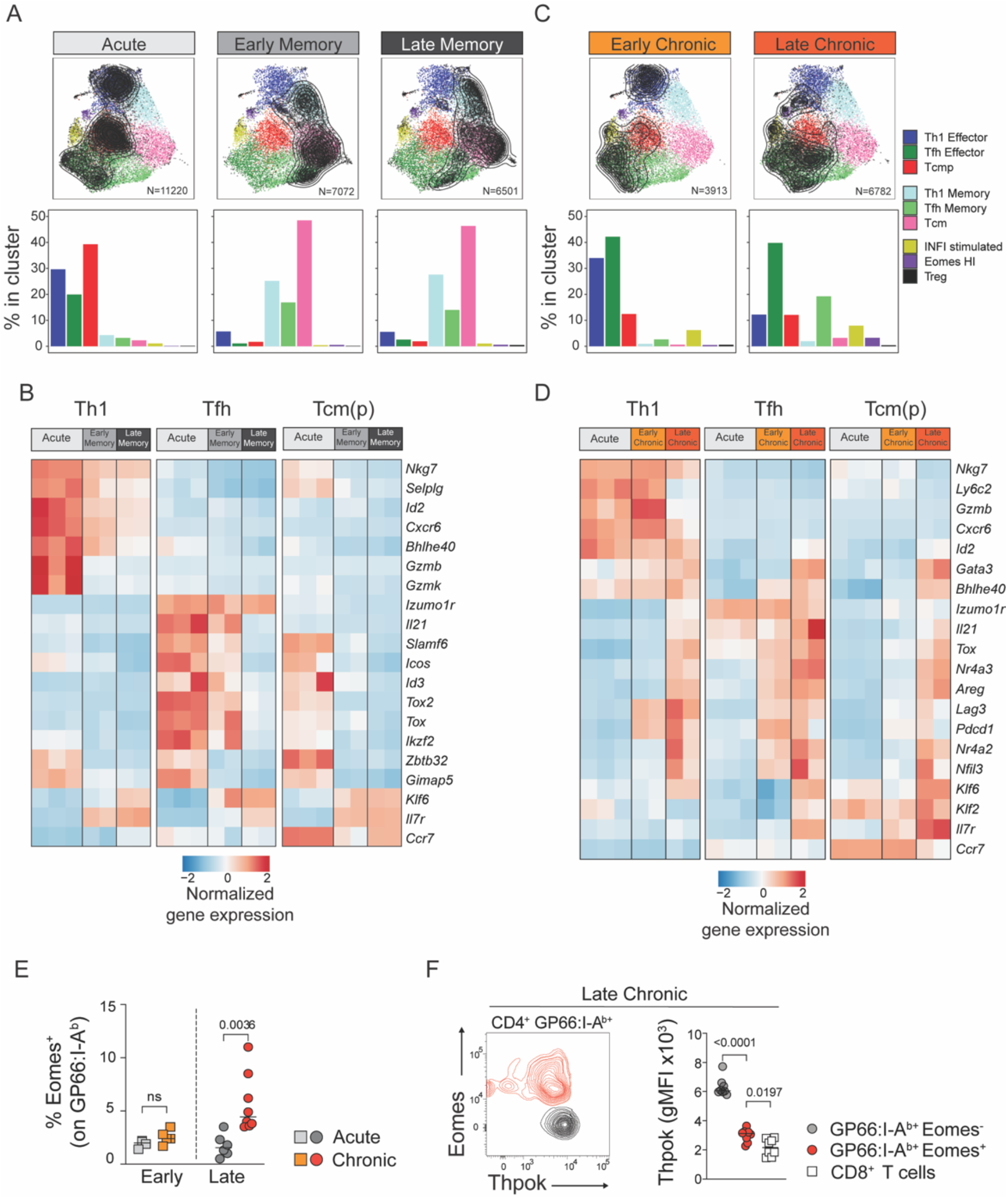
Subset-specific adaptation of CD4^+^ T cell states in chronic and acute infections. **A,C)** Distribution of T cell states at different timepoints after acute and chronic infections. In the UMAP plots, contour lines indicate the density of T cells for each type of infection and timepoint; the barplots in the bottom row indicate the percentage of cells in each subtype in the indicated condition. **B,D)** Normalized average expression during acute and chronic infections among Th1, Tfh and Tcm(p) subtypes. Selected genes from differentially expressed genes shown in Fig. S3A,C. **E,F)** Animal were infected with LCMV Armstrong (Acute) or Clone 13 (Chronic) and analyzed at the indicated timepoints (Early: 7dpi; Late: 21dpi). **E)** Graph shows the percentage of Eomes^+^ cells among spleen GP66:I-A^b+^T cells. **F)** Plot (left) shows the intracellular expression of Eomes and Thpok on spleen GP66:I-A^b+^T cells analyzed 21 dpi after LCMV Clone 13 infection. Graph (right) shows Thpok mean fluorescent intensity (gMFI) in the indicated population. E,F are from one experiment with >5 mice per group, representative of 2 independent experiments.

To further investigate the transcriptional changes underlying this transition, we sought to identify gene expression differences among matching states during the acute response and early memory phase. We also interrogated potential changes between the early and late memory timepoints resulting in the identification of 183 genes differentially expressed in a time- and state-specific manner in response to acute infection **(*Fig. S3A*, *Supplemental Data 2*)**. Our analyses revealed that the transition of Th1 and Tfh subtypes from effector phase to memory phase were accompanied by the dampening of effector function molecules such as *Gzmb* (Th1) and *Il21* (Tfh), and by the acquisition of *Il7r* expression at the memory phase **(*Fig 3B*)**. However, the downregulation of effector programs, especially for the Tfh state, was more pronounced in late memory phase compared to early memory, suggesting that memory CD4^+^ T cells undergo continued transcriptional remodeling after the contraction phase. In contrast, the central memory-type cells [Tcmp and Tcm clusters, referred to as Tcm(p)] readily downregulated most effector-associated genes at the early memory phase. Similarly, the Tcm(p) state more quickly upregulated genes associated with the function, survival or trafficking of memory cells like *Ccr7*, *Il7r*, *Bcl2* or *Klf6* compared to Th1 and Tfh states. This early divergence and stable expression of memory genes is compatible with the idea that Tcmp represent a pool of circulatory cells with an increased fitness to develop into long-lived memory cells.

These analyses highlight subtype-specific transcriptional changes from effector to memory states. In particular, they suggest that, while Tcmp are poised to transition to memory, Th1 and Tfh states do so with different kinetics and using divergent transcriptional modules.

### Subtype-specific adaptation of CD4^+^ T cell states to chronic infection

We next investigated the adaptation of CD4^+^ T cell transcriptional states to chronic infection. Compared to the acute setting, we did not observe a sharp transition to memory states at day 21 and most virus-specific CD4^+^ T cells matched effector subtypes **(*Fig. 3C*)**. Although the proportion of the Th1 subtype remained similar at early timepoints between acute and chronic condition, there was a reduction in the Th1 effector subsets at late chronic stages. In addition, we observed a larger pool of Tfh cells both at early and late chronic timepoints compared to acute settings **(*Fig. 3C*)**, consistent with previous studies highlighting a Tfh bias during Clone 13 infection (*Brooks et al., 2005; Fahey et al., 2011*). We also noted that the fraction of Tcmp cells was lower in chronic infection, both at early and late stages, compared to the CD4^+^ T cells in acute infection. Indeed, the proportion of Tcmp and Tcm cells, which can be identified by flow cytometry based on CCR7 expression, was greatly reduced among virus-specific CD4^+^T cells in chronic settings compared to acute infection **(*Fig. S3B*)**.

Next, we aimed to assess the transcriptional changes affecting the differentiation and persistence of each subtype in responses to chronicity. To this end, we measured the differences across subtypes at an early (7 dpi) and late phase (21 dpi) of the chronic response. We identified 164 genes differentially expressed between Acute vs. Early Chronic or Early vs. Late Chronic timepoints **(*Fig. S3C, Supplemental Data 2*)**. Importantly, most changes observed at the late chronic phase were not present at the early chronic stage, suggesting that they are not merely attributed to changes in viral replication or host responses to viral variants. We observed that late chronicity was associated with the upregulation of a shared gene module, including Nr4a family members (*Nr4a1*, *Nr4a2*, *Nr4a3*) and *Tox* in all subtypes **(*Fig. 3D, S3C*),** indicative of the strong TCR engagement in response to persistent antigen (*Seo et al., 2019).* Similarly, the expression of inhibitory receptors such as *Pdcd1* (encoding PD-1) and *Lag3* was also detected across states in late chronic samples. Late timepoints were also characterized by the downregulation of effector modules, including gene associated with cytotoxic function, such as *Gzmb* and *Ctsw (encoding Cathepsin W)* in Th1 clusters. In contrast, the expression of *Il21*, as well as the transcription factor *Maf* remained highly expressed in Tfh clusters suggesting that, unlike in Th1, effector functions in Tfh cells are not dampened at late chronic phase. In fact, *Il21* expression appears to increase at late timepoints in both Tfh and Th1 subtypes **(*Fig. 3D*)**. In contrast to what was observed in response to acute infection, Tcm(p) minimally diverged from other states, as the expression of inhibitory receptors and transcription factors associated with T cell dysfunction such as *Ikzf2* (encoding Helios) *Bhlhe40* or *Gata3 (Crawford et al., 2014; Doering et al., 2012; Singer et al., 2016)* were equally upregulated in all states in chronic settings*. However,* unlike other subtypes, Tcm(p) maintained expression of *Il7r*, *Ccr7* and *S1rp1* (***Fig. 3D*, *S3C***).

In addition to the 6 main CD4^+^ T cell states, we detected two distinct states that were almost exclusively present in response to chronic infection: the IFNI-stimulated state and the *Eomes*-HI state **(*Fig. 3AC*)**. The *Eomes*-HI state was specifically observed at late stages of chronic infection, consistent with previous studies (*Crawford et al., 2014; Lewis et al., 2016*) and flow cytometry analyses (***Fig. 3E and S3D***). Because this cluster was characterized by the co-expression of *Eomes*, *Lag3 and Xcl1* (***Fig. S3E***), three functional targets repressed by the CD4 T cell-defining transcription factor Thpok (*Ciucci et al., 2019; Taniuchi, 2018*), we sought to determine whether its expression was altered in this subset. Indeed, we found that Thpok protein expression was reduced specifically in Eomes^+^ virus-specific CD4^+^ T cells late during chronic infection (***Fig. 3F***).

In summary, these analyses showed that persistent antigen exposure during chronic infections deeply alters CD4^+^ T cell differentiation by imprinting both common and subtype-specific transcriptional changes associated with chronicity.

### Clonotype-fate relationships of virus-specific CD4^+^ T cells in acute and chronic infections

Because cell-intrinsic factors, notably the expression of TCR, have been shown to impact the differentiation of virus-specific T cells (*Khatun et al., 2021; Kunzli et al., 2021; Snook et al., 2018*), we sought to determine whether CD4^+^ T cell states are influenced by the expression of particular sets of TCR chains. To describe the clonal relationship between CD4^+^ T cell states we analyzed the TCR usage of all T cells for which a productive pair of TCRα/TCRβ sequences was detected (65% of all single cells). To limit potential confounding factors related to the survival or expansion of clonotypes over time, we restricted our analyses to early timepoints (*i.e.* 7 dpi) after acute or chronic infections. We observed a consistent pattern of clonal expansion across different animals, with large clonotypes (>20 cells) occupying roughly half of the clonal space in all samples, both for acute and early chronic settings ***Fig. 4A***). Next, we interrogated potential repertoire overlaps between animals considering the pair of nucleotide and amino-acid sequences of the CDR3 regions. Strikingly, this analysis showed that less than 3% of the clonotypes (7 out of 795 CDR3 nucleotide pairs; 20 out of 779 CDR3 protein pairs) were observed in two or more animals, when considering clones with 3 cells or more (***Fig. 4B***). Similar results we obtained when all clones, including singletons, were included (***Fig. S4A***). This suggests that the TCR repertoire of virus-specific CD4^+^ T cells is largely private, *i.e.* subject-specific, even when considering a single epitope and animals with the same genetic background.

**Figure 4:**
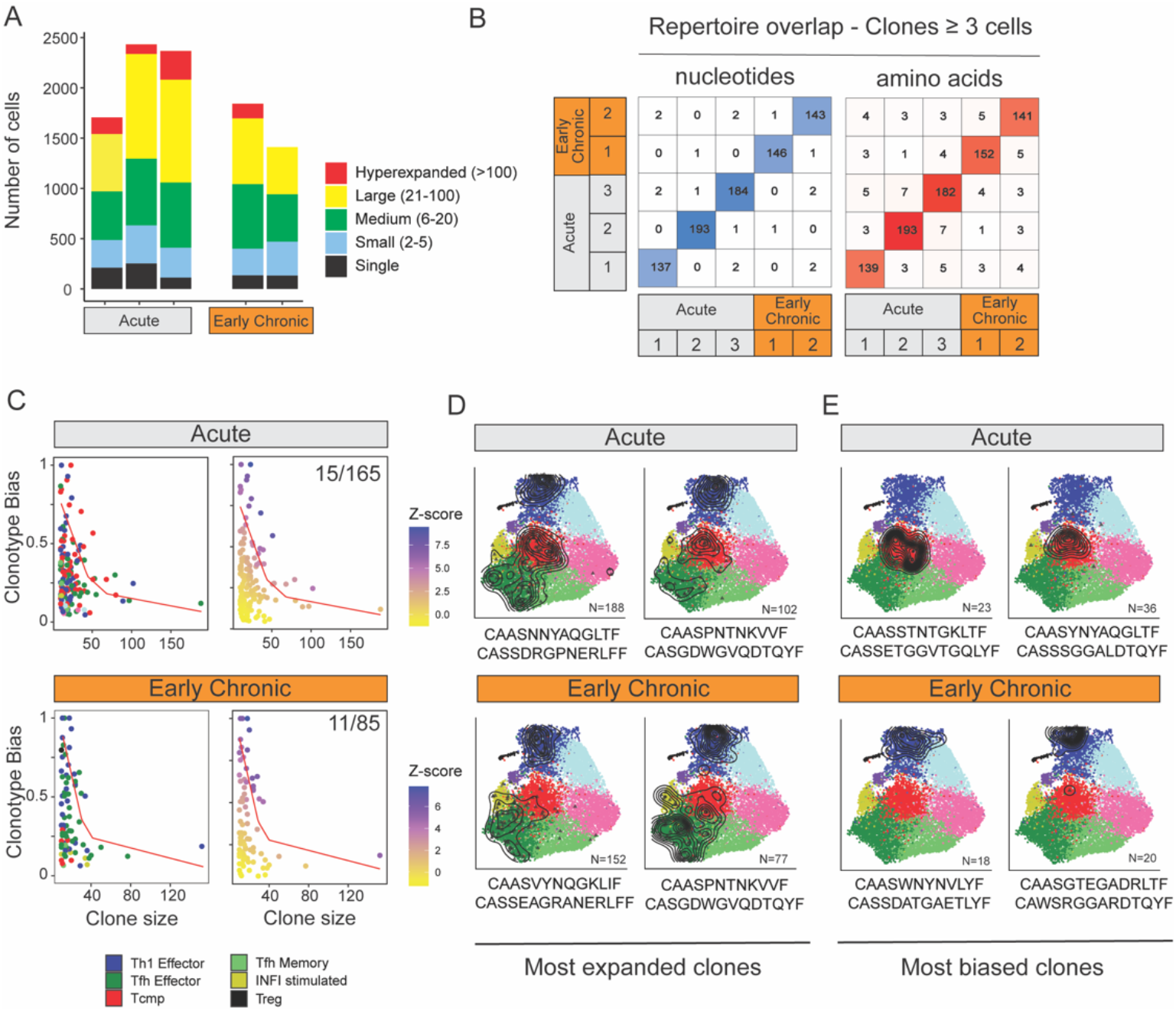
Clonal structure of virus-specific CD4^+^ T cells and clonotype-fate relationship. **A)** Fraction of clonal space occupied by clonotypes with different levels of expansion. For each sample (three replicates for Acute, two replicates for Early Chronic), the plot indicates the number of single cells that belong to clonotypes in one of the five classes of expansion. **B)** Number of clones with identical CDR3 nucleotides or amino acid sequence pairs between individual samples among clonotypes with ≥ 3 cells. **C)** Clonotype bias analysis for acute and chronic infection samples. Plots show clonotype bias vs. clonal size for all clones with > 10 cells. Clonotypes are colored by predominant T cell subtype (left) or by Z-score of the clonotype functional bias (right). Red line highlights the null distribution (background distribution) for each condition. **D, E)** Distribution of cells over the reference atlas for the D) four most expanded and E) four most biased clonotypes in acute and chronic infections. The CDR3 alpha and beta sequences and the clonotype size are indicated for each clonotype.

Next, we explored the potential relationship between TCR usage and the emergence of specific T cell states at early infection timepoints. To this end, we defined a “clonotype bias” metric to quantify how individual clones are skewed towards a specific subtype (see methods). At its extreme values, a clonotype bias of 1 indicates that a clonotype is composed uniquely of cells from the same subtype, and a clonotype bias of zero corresponds to a clonotype that matches exactly the background subtype distribution of the whole sample. Because small clonotypes are statistically more likely to show high clonotype bias compared to large clonotypes, we only considered expanded clonotypes with 10 or more cells, and corrected for clonotype size by generating expected background distributions by random permutation (***Fig. S4B***, see methods). This analysis revealed that most clonotypes were largely unbiased, both in acute and chronic infections, giving rise to multiple CD4^+^ T cell states (***Fig. 4C-D***), consistent with previous studies in acute infection (*Khatun et al., 2021*). However, we observed that a small number of clonotypes exhibited significant functional bias (Z-score>5), i.e. they were preferentially enriched in one specific subtype (***Fig. 4C***). In the case of acute infection, 9% of expanded clonotypes (15 out of 165) exhibited a functional bias toward one subtype. Similarly, 13% of expanded clonotypes in chronic infection (11 out of 85) showed a significant clonotype bias (***Fig. 4C***). Interestingly, in acute infection 14 out of 15 biased clonotypes were skewed toward either the Tcmp or Th1 states (8 and 6 respectively), while in chronic infection all 11 biased clonotypes showed functional bias towards either Th1 or Tfh states (8 and 3 respectively) (***Fig. 4C-E***).

These combined analyses of the TCR repertoire and transcriptional landscape reveals that the vast majority of the clonotypes can differentiate into multiple states with minimal functional skewing. However, a minority of these clonotypes are significantly biased towards a particular functional state, and this bias appears to be influenced by the type of infection.

### Interpreting CD4^+^ T cell states across studies and conditions using the new reference atlas

A cell atlas is particularly useful when it serves as a “reference” to compare and interpret new data. We have recently proposed a computational method, ProjecTILs, that allows analyzing single-cell datasets by projection into a reference atlas (*Andreatta et al., 2021*). Using this approach, we sought to apply our new CD4^+^T cell reference atlas for the interpretation of CD4^+^ T cell differentiation in external datasets from several different studies (***Fig. 5A***).

**Figure 5:**
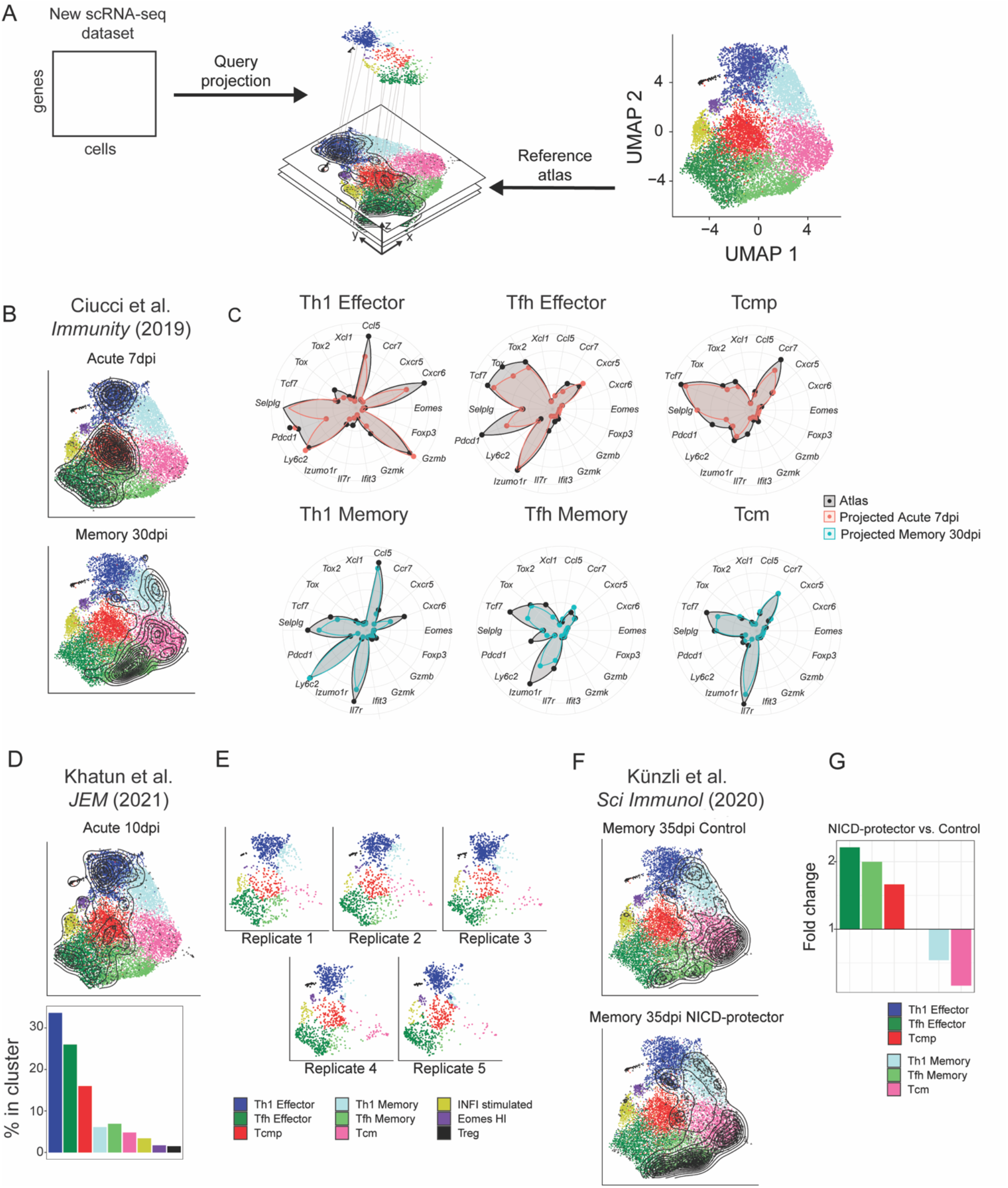
Interpretation of CD4^+^T cell states in external datasets by projection into the reference atlas. **A)** New scRNA-seq datasets from additional studies can be mapped to the CD4^+^T cell reference atlas using the ProjecTILs algorithm and interpreted in the context of the space and cell states of the reference. **B)** UMAP embeddings for the data by Ciucci et al. at 7 and 30 days after acute LCMV infection. **C)** Expression profiles for a panel of marker genes for the projected cells from Ciucci et al. (blue and red) compared to the reference atlas profiles (black). **D)** UMAP embeddings and subtype composition for the virus-specific CD4^+^ T cell scRNA-seq data by Khatun et al. 10 days after LCMV acute infection. **E)** Distribution of cells for five individual replicates from the data by Khatun et al. **F)** UMAP embeddings for the virus-specific CD4^+^T cell scRNA-seq data by Kunzli et al. 35 days after LCMV acute infection, for NAD-induced cell death (NICD)-protector treated mice and control. **G)** Fold-change for composition of CD4^+^T cells states in NICD-protector vs. control sample, highlighting relative increase and decrease in the frequency of individual cell subtypes. Black contour lines in all UMAP identify the density of single cells from the projected dataset, overlaid on the space of the reference

To verify the accuracy of dataset projection into our reference atlas, we re-analyzed data comprised of scRNA-seq measurements of LCMV-specific CD4^+^ T cells isolated at 7 and 30 days post-infection with LCMV Armstrong (*Ciucci et al., 2019*). Validating both the accuracy of the atlas and of the projection algorithm, cells from day 7 were projected into the Tcmp and effector states, while cells from day 30 were largely projected into the memory states (***Fig. 5B***). Importantly, the expression profile of key marker genes for the projected samples matched closely to the expression profile of the reference atlas in all major cell subtypes, confirming the accuracy of external data projection (***Fig. 5C***).

Seeking to further explore temporal differences in acute infection, we re-analyzed LCMV-specific CD4^+^ T cells isolated 10 days post-infection with LCMV Armstrong (*Khatun et al., 2021*). Dataset projection into our CD4^+^ T cell reference atlas revealed that the majority of these virus-specific T cells were found in the Th1 Effector, Tfh Effector or Tcmp states (***Fig. 5D***), similarly to subtype compositions we observed at day 7 in acute infection (***Fig. 3A***). This is consistent with the notion that transition to memory phenotypes occurs later, at day 12-20 post-infection (*Marshall et al., 2011*). Importantly, a very similar subtype distribution was observed in different mice, highlighting the robustness of the projection algorithm across multiple biological replicates (***Fig. 5E***).

Finally, we tested the robustness of our projection approach in detecting variations across biologically similar samples. To this end, we took advantage of transcriptomic datasets of LCMV-specific CD4^+^ T cells isolated at memory timepoints (day 35) after in vivo administration of an inhibitor blocking NAD-induced cell death (NICD) (*Kunzli et al., 2020*). As expected for a late timepoint, in both the control and treated conditions most cells were projected into the memory clusters (Tcm, Th1 memory and Tfh memory) (***Fig. 5F***). However, the NICD-protector-treated sample showed a ~2-fold increase of Tfh effector and Tfh memory cells compared to control (***Fig. 5F-G***). This is consistent with the hypothesis that Tfh cells are more susceptible to NICD than other CD4^+^ T cell subtypes, and that *in vivo* NICD-blockade can enhance the persistence of Tfh populations after infection (*Kunzli et al., 2020*). Thus, these analyses show that our atlas and projection method are able to capture even subtle alterations in the distribution of transcriptional states across experimental conditions.

The virus-specific CD4^+^ T cell reference atlas developed in this study can be explored within the SPICA portal at https://spica.unil.ch/refs/viral-CD4-T, where users can compare the expression of genes of interest in individual cell subtypes and across the reference space. SPICA also hosts interactive analyses for the datasets described above and for many others (https://spica.unil.ch/projects). Finally, researchers can project their own scRNA-seq data through the SPICA web interface, or by using our R package ProjecTILs available at https://github.com/carmonalab/ProjecTILs.

## Discussion

CD4^+^ T cells orchestrate immune responses to pathogens and critically support protection conferred by vaccination. However, the phenotypic and functional plasticity of CD4^+^ T cells has hindered a robust, unbiased delineation of pathogen-specific T cell subtypes. Although the precise characterization of T cell transcriptional states is fundamental towards understanding the dynamics of immune responses, the subtype-specific changes occurring over time in response to acute and chronic infections remain poorly understood.

In this work, we aimed at identifying the transcriptional and clonal landscape of polyclonal antigenspecific CD4^+^ T cells in acute and chronic infections by single-cell transcriptomics. By combining robust animal models, efficient bioinformatics algorithms for data integration and expert annotation, we provide a reference atlas of virus-specific CD4^+^ T cells. Integrated in a computational framework for reference projection, this atlas enables interpretation of new, external T cell dataset, providing a powerful resource for the community.

We provide new insight into the transcriptional adaptation of virus-specific CD4^+^ T cell populations over time and across conditions. Our analyses during acute infection highlight that, although the major gene expression changes occur at the end of the initial proliferative burst, early memory CD4^+^ T cells that survive the contraction phase undergo continued transcriptional remodeling at later timepoints, similarly to late changes at play in CD8^+^ and NK T cells memory development (*Chang et al., 2014; Lau et al., 2018; Milner et al., 2020*). However, CD4^+^ T cell subsets undergo memory transition in a divergent manner, where each subtype acquires memory features with different kinetics and using non-overlapping transcriptional modules. Similar to the acquisition of the memory program in CD8^+^ T cells, Th1 memory differentiation is characterized by a dampening of effector functions accompanied by the upregulation of molecules associated with their long-term survival. In contrast, the transition of effector Tfh cells to memory states appears to be delayed, as molecules associated with Tfh function such as IL-21 or ICOS remain expressed in early memory Tfh cells, and their expression only decreases at later timepoints. This delayed transition into resting memory could be associated with differential and prolonged antigen exposure of Tfh within the germinal centers compared to Th1 cells (*Kunzli et al., 2020*). However, it is interesting to note that, even at later timepoints, Tfh cells do not acquired the same memory program than Th1 cells. This observation is consistent with the idea that Tfh cells diverge early from other CD4^+^T functional subsets and transition to memory using transcriptional modules unique to this state (*Ciucci et al., 2019; Hale et al., 2013*). Interestingly, the transition to the Tcm state diverge from both Th1 and Tfh. In fact, most memory-associated features like the expression of *Ccr7, Il7r* or *Bcl2* appear quickly at the early memory phase or are already present in Tcmp cells at the acute phase of the response. This observation is in line with the concept that Tcmp cells already express a large fraction of memory-associated genes allowing for their survival and homing to lymphoid organs. Thus, it is possible that most Tcm derived mainly from Tcmp cells through the acquisition of a transcriptional state poised for memory differentiation, similar to memory-precursors in CD8^+^ T cell populations (*Joshi et al., 2007; Kaech et al., 2003*). In any event, these developmental relations remain speculative in the absence of robust lineage tracing data.

During chronic infection, similar state-specific changes occur in CD4^+^ T cell subtypes. In fact, in addition to a shared transcriptional module upregulated in all subsets, Th1 and Tfh states largely differ in their adaptation to chronic antigen stimulation typically associated with dysfunction. Imprinting of persistent antigen exposure on Th1 cells results in a reduction of effector function characterized by the repression of effector molecules like granzymes, reminiscent to CD8^+^ T cell function dampening in chronic infection and cancer (*Crawford et al., 2014; Singer et al., 2016*). In contrast, Tfh effector functions remain unaffected at the chronic phase of the response. While this could be related to the strong Tfh bias observed in chronic infections (*Fahey et al., 2011*), we surprisingly observed in Th1 subsets features typically associated with Tfh cells, including the expression of IL-21. Because IL-21 is critical to limit T cell dysfunction during chronic infections (*Elsaesser et al., 2009; Frohlich et al., 2009; Yi et al., 2009*), it is possible that compensatory mechanisms enforce its expression in non-Tfh subsets. Alternatively, the “boundaries” between CD4^+^ T cells states may be less easily delineated during chronic infection. In line with this idea, our analyses reveal that during chronic infection, the accumulation of Eomes^+^virus-specific CD4^+^ T cells is accompanied by the downregulation of the CD4 T cell-defining factor Thpok. Similar to “redirected” CD4^+^ T cells in both human and mouse (*Mucida et al., 2013; Serroukh et al., 2018*), this CD4^+^ T cell subset is characterized by the upregulation of targets actively repressed by Thpok, including the transcription factor Eomes and genes associated with cytotoxic activity (*Ciucci et al., 2019; Vacchio et al., 2019*). While the downregulation of Thpok in CD4^+^T cells can be influenced by the cytokine milieu (*Cervantes-Barragan et al., 2017; Reis et al., 2014*), it is important to note that the Eomes^+^ Thpok^low^ CD4^+^ T cell population appears to be limited to chronic settings, including in responses to the gut microbiota (*Cervantes-Barragan et al., 2017; Mucida et al., 2013*). Thus, it is possible that both environmental cues and chronic antigen activation either at mucosal surface or lymphoid organs are required for the development of this subset.

This study represents the first step into building a more comprehensive map of the transcriptional landscape supporting the functional heterogeneity of pathogen-specific CD4^+^ T cells. Future iterations will integrate additional conditions such as virus-specific CD4^+^ T cells from non-lymphoid tissues to identify tissuespecific adaptation of CD4^+^ T cell states in both mouse and human. We provide a powerful and user-friendly resource for investigators to explore the new CD4^+^ T cell atlas and to interpret their own datasets in the context of this reference.

## Supporting information

DataSupplemental1

DataSupplemental2

DataSupplemental3

---

**Figure Supplemental 1:**
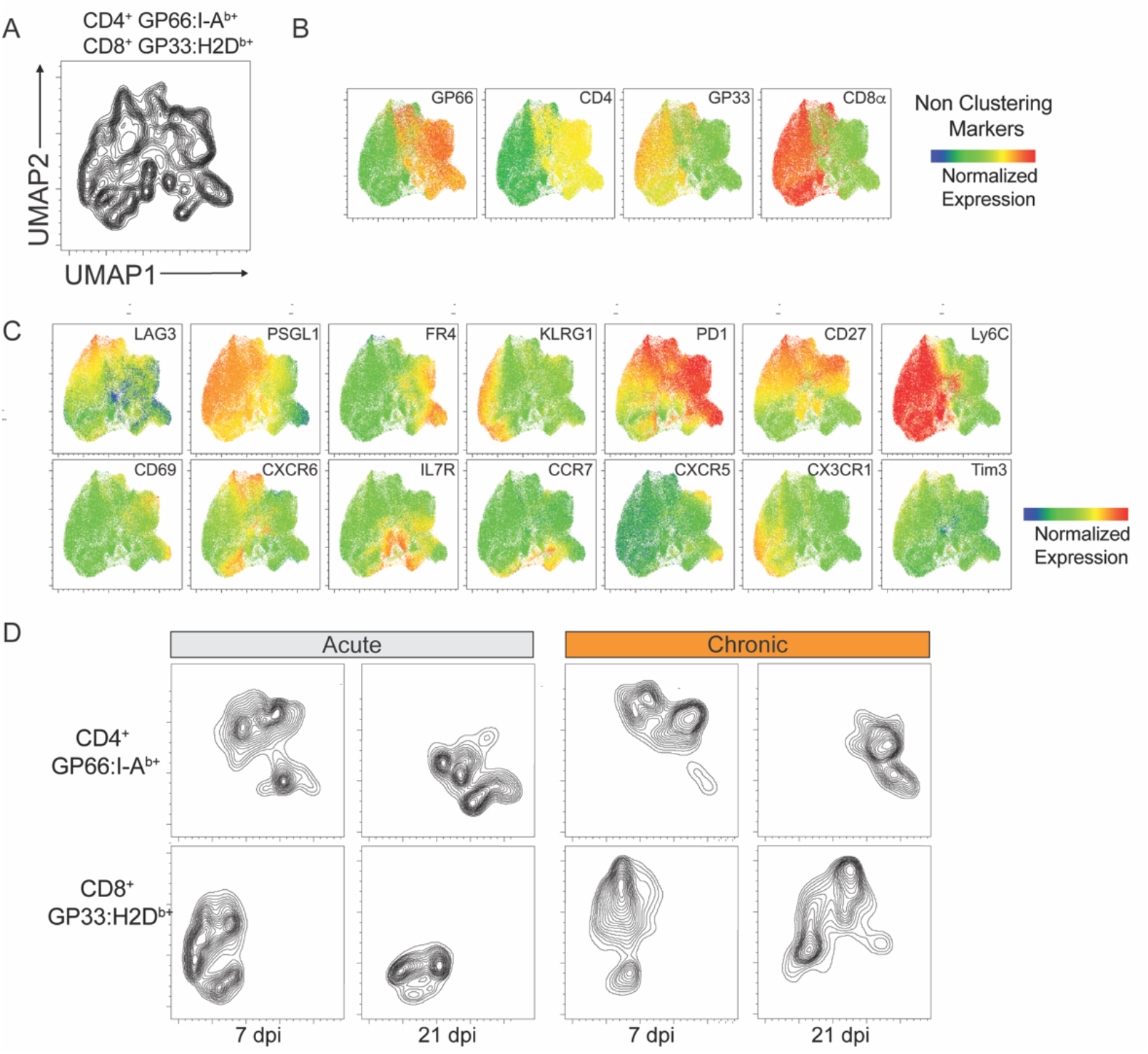
Phenotypic characterization of virus-specific T cells. GP66:I-A^b+^ CD4^+^ and GP33:H2D^b+^ T cells were analyzed 7 and 21 dpi after infection with LCMV Armstrong and Clone 13. **A)** UMAP was calculated based on the expression of 14 markers on virus-specific T cells. **B)** CD4, CD8 and tetramer levels shown on the UMAP as in A). **C)** Expression of 14 parameters used to generate the UMAP. **D)** T cells from the indicated conditions were projected on the UMAP. Experiment with 4-5 mice per group, representative of 2 independent experiments.

**Figure Supplemental 2:**
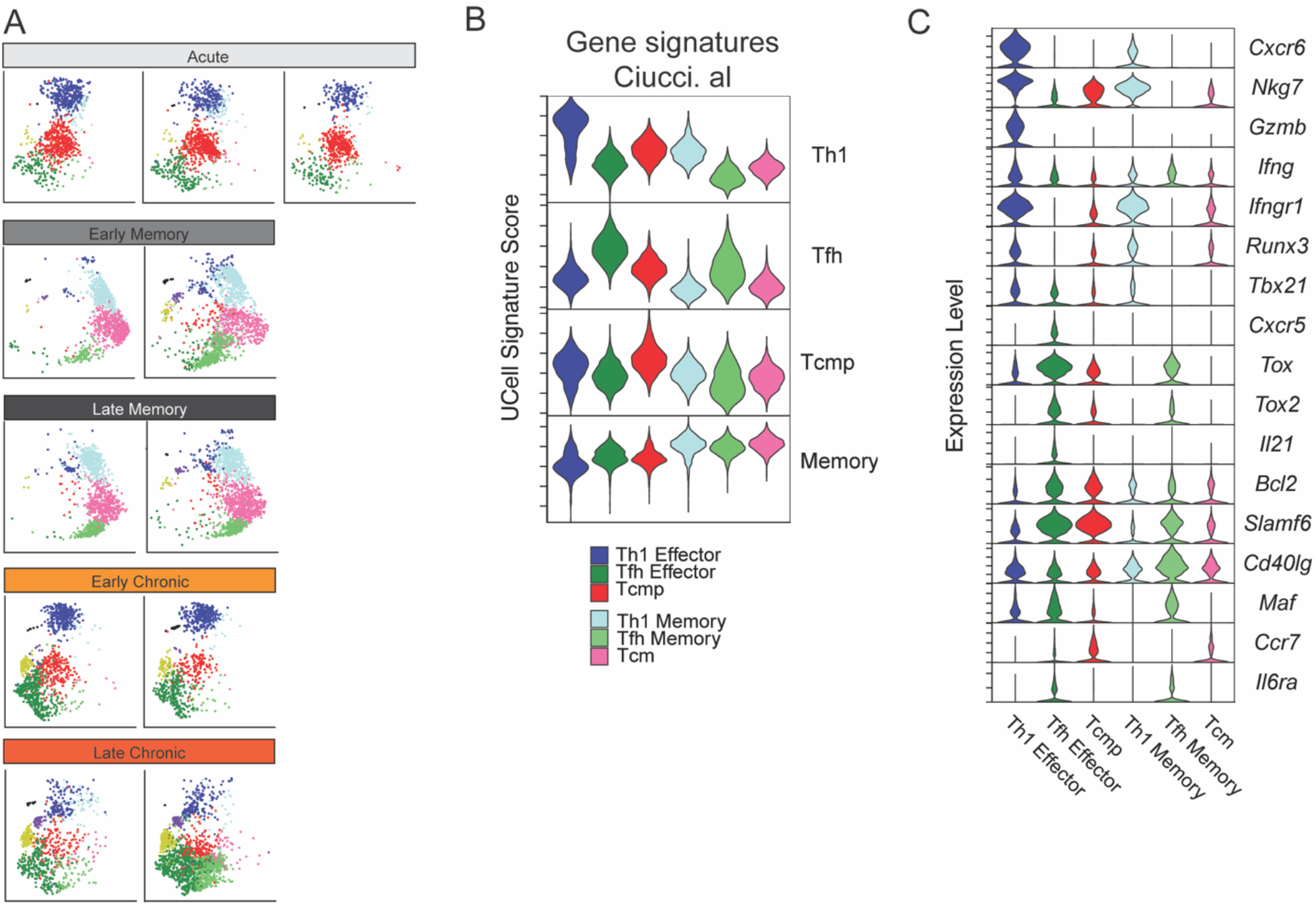
Distribution and signatures of CD4^+^ T cell states during infections. **A)** Distribution of individual samples in the UMAP space of the reference atlas as described in Fig. 2. Samples are grouped by timepoint and condition. **B)** Relative expression of gene signatures as defined in Ciucci et al. calculated using UCell and **C)** Relative gene expression of selected genes on cells assigned to six main CD4^+^ T cell states.

**Figure Supplemental 3:**
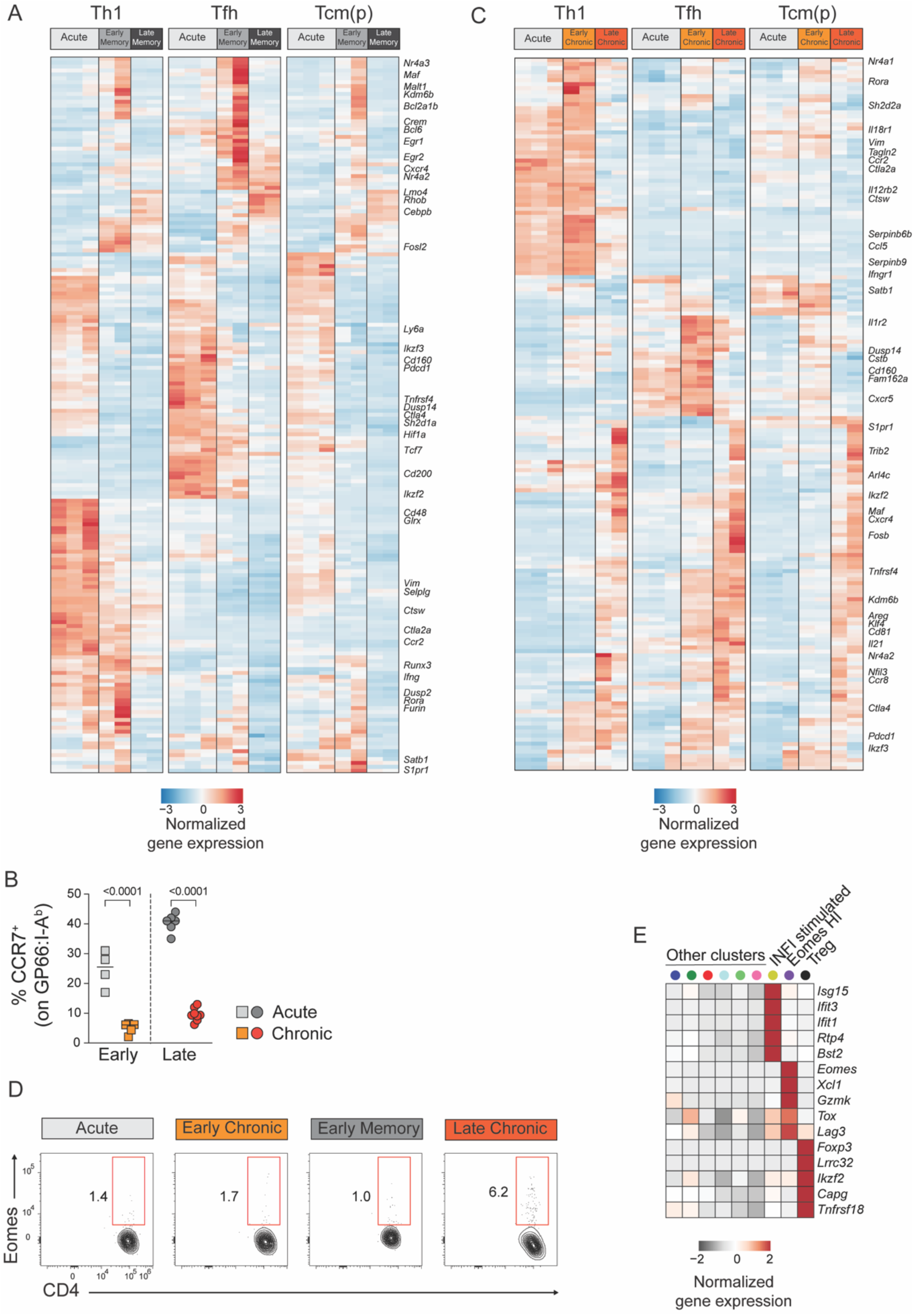
Adaptation of CD4^+^ T cell states in chronic and acute infections. **A,C)** Normalized average expression during acute and chronic LCMV infections among Th1, Tfh and Tcm(p) subtypes. **B,D)** Animal were infected with LCMV Armstrong (Acute) or Clone 13 (Chronic) and spleen GP66:I-A^b+^T were analyzed at the indicated timepoints (Early: 7dpi; Late: 21dpi). **B)** Graph shows the percentage of CCR7^+^ cells and **D)** the expression of Eomes and CD4 at the indicated timepoints. Date are from one experiment with >5 mice per group, representative of 2 independent experiments. **E)** Average expression of differentially expressed genes in the 3 minor subtypes of the reference atlas.

**Figure Supplemental 4:**
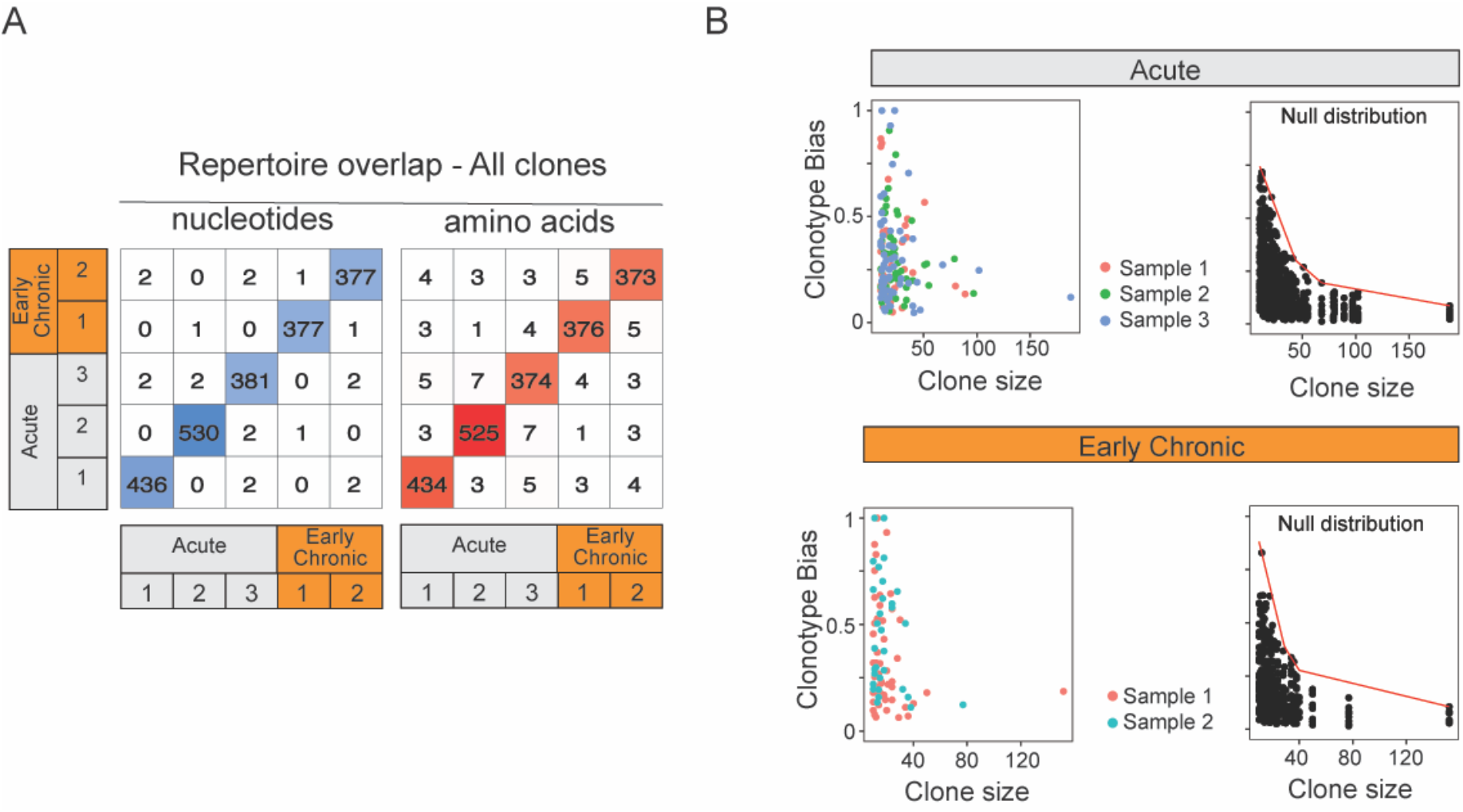
Repertoire overlap of virus-specific CD4^+^ T cells and clonotype bias. **A)** Number of clones with identical CD3 nucleotides or amino acid sequence pairs between individual samples among all clonotypes. **B)** Clonotype bias analysis for acute and chronic infection samples. A clonotype bias of 1 indicates that a clonotype is composed uniquely of cells from the same subtype, and a clonotype bias of zero corresponds to a clonotype that matches exactly the (random) background subtype distribution. Left plots show clonotype bias colored by sample for all clone with > 10 cells. Left plot shows null distribution (background distribution) for each condition.

## Methods

### Mice, virus and infections

C57BL/6Ncr were infected by intra-peritoneal injection of 2 x 10^5^ pfu of LCMV Armstrong or intravenously with 2 x 10^6^ pfu of LCMV Clone 13. Viral stocks were prepared and titrated as previously described (*Ciucci et al., 2019; Dangi et al., 2020*).

### Antibodies

Antibodies for the following specificities were purchased either from Becton-Dickinson Pharmingen, BioLegend or ThermoFisher: CD4 (GK1.5), CD8α (53-6-7), CD5 (53-7.3), B220 (RA3-6B2), CD44 (IM7), IL-7Ra (A7R34), CCR7 (4B12), CXCR5 (SPRCL5), CXCR6 (SA051D1), PSGL1 (2PH1), Ly6C (HK1.4), CD27 (LG/3A10), FR4 (12A5), Thpok (T43-94), Eomes (Dan11mag), CD69 (H1.2F3), LAG3 (C9B7W), Tim3 (RMT3-23), KLRG1 (2F1), PD1 (29F.1A12), CX3CR1 (SA011F11). MHC tetramers loaded with the LCMV GP66 or GP33 peptides were obtained from the NIH Tetramer Core Facility.

### Cell preparation and staining

Spleen cells were prepared and stained as previously described (*Ciucci et al., 2019*). Surface staining with GP66:I-Ab tetramer and for CCR7 or CXCR5 was performed at 37°C for 1 hour prior to staining with antibodies for other cell surface markers. Intracellular stainings were performed as previously described using the Transcription Factor Staining Buffer (ThermoFisher) (*Chopp et al., 2020*). Data was acquired on Aurora spectral flow cytometer (Cytek) and analyzed with FlowJo software (TreeStar). Dead cells and doublets were excluded by DAPI or LiveDead staining (Invitrogen) and forward scatter height by width gating. Purification of lymphocytes by cell sorting was performed on a FACS Fusion and FACS Aria (BD Biosciences).

### Single-cell RNA sequencing

GP66:I-A^b+^ spleen T cells were sorted from LCMV infected animals, loaded onto the Chromium platform (10X Genomics) to generate cDNAs carrying cell- and transcript-specific barcodes that were used to construct sequencing libraries using the Chromium Single Cell 5’ or 3’ Library & Gel Bead Kit according to the manufacturer instructions. For pooled captures, 2 cell populations were sorted and barcoded separately with TotalSeq antibodies (BioLegend) before mixing and cell captures **(*Supplemental Data 1*)**. Libraries were sequenced on multiple runs of Illumina NextSeq or Novaseq using paired-end reads to reach a sequencing saturation of 60-90%, resulting in at least 2-9×10^4^ reads/cell. Single-cell sequencing files were processed, and count matrixes extracted using the Cell Ranger Single Cell Software Suite (10X Genomics).

### scRNA-seq data processing and quality control

Single-cell transcriptomes and single-cell TCR sequences were mapped and combined using the *combineTCR* function from scRepertoire (*Borcherding et al., 2020*). We performed quality control on the single-cell data using the following criteria: number of detected genes > 700; number of UMIs > 1500 and < 15,000; percentage of ribosomal genes < 50 and percentage of mitochondrial genes < 10. For all these parameters, we additionally removed all extreme outlier cells outside the 1st and 99th percentile in each sample. In order to filter out potential contaminants and experimental artifacts, we applied the UCell package (*Andreatta and Carmona, 2021b*) to evaluate a panel of signatures for several common immune and non-immune cell types. This resulted in high quality transcriptomes for 35,488 single cells from 11 samples, covering acute and chronic infections at three different timepoints.

### scRNA-seq data integration

For the construction of the CD4^+^ T cell reference atlas, datasets were downsampled to balance the contribution from different types of infection and timepoint. To this end, a maximum of 5,000 cells were randomly selected for each of the 5 subsets: acute day 7, acute day 21, acute day 60, chronic day 7 and chronic day 21. To mitigate batch effects between samples, we integrated the 11 samples using STACAS (*Andreatta and Carmona, 2021a*) with the following parameters: number of variable genes=800, dist.thr=0.6, dims=20. On the integrated data, unsupervised clusters were calculated using the *FindNeighbors* and *FindClusters* functions from Seurat (*Hao et al., 2021*) with parameters: k.parameter=10, resolution=0.4. Finally, unsupervised clusters were manually annotated guided by differential expression analysis between clusters, merging clusters where appropriate, to obtain nine “functional clusters” that summarize the diversity of CD4^+^ T cells in acute and chronic infections.

### Reference atlas projection of scRNA-seq data

In order to avoid large imbalances between subtypes, and to limit its disk size, the CD4^+^ T cells reference atlas is a downsampled version of all available data. Thus, low-dimensional embeddings and subtype annotations for all cells generated in this study were obtained by dataset projection with ProjecTILs using default parameters (*Andreatta et al., 2021*). Similarly, the same method was applied to project and interpret scRNA-seq data from three additional studies not included in the reference map (*Ciucci et al., 2019; Khatun et al., 2021; Kunzli et al., 2020*).

### Gene signatures of adaptation to acute and chronic infections

CD4^+^ T cell state signatures were calculated by comparing each of the 6 main clusters to the two other clusters in the same state (i.e. Effector or Memory) using the *FindMarkers* function implemented in Seurat. To summarize subtype-specific transcriptional changes at different stages of infection, we first identified differentially expressed genes between timepoints and subtypes. For this analysis, Th1 effector and Th1 memory cells were grouped together to identify Th1-type cells. Similarly, we combined the Tfh effector and Tfh memory clusters (Tfh type), as well as Tcmp and Tcm clusters (Tcm(p) type). For significantly differentially expressed genes, we calculated average expression profiles for the Th1, Tfh and Tcm(p) types at individual timepoints using the *find.discriminant.genes* function of ProjecTILs (*Andreatta et al., 2021*). The UCell algorithm (*Andreatta and Carmona, 2021b*) was applied on the CD4^+^ T cell atlas subtypes to evaluate gene signatures for subtypes identified in a previous study (*Ciucci et al., 2019*). To excluded potential confounding factors, ribosomal-associated, sex-specific and as TCR induced transcripts (*Magen et al., 2019*) were removed from signatures related to acute and chronic adaptation **(*Supplemental Data 3*)**.

### T cell clonal analysis

The CDR3 amino acid sequence for productive alpha-beta VDJ rearrangements obtained by scTCR-seq were used as unique “barcodes” to identify individual T cell clones. The expansion level of a clone was calculated as the absolute number of cells with identical TCR sequence in a given sample, either in terms of CDR3 sequence or full nucleotide sequence. Expanded clones that were unique to a sample were denoted as private clones, expanded clones found in at least three samples were denoted as public clones. Merging and visualization of scTCR-seq data were performed using the scRepertoire package (*Borcherding et al., 2020*).

To measure gene expression-TCR relationship, we defined a “clonotype bias” metric to quantify whether a given clonotype was preferentially composed of one of the T cell subtypes: c = max_i_ [ (f_i_ - q_i_) / (1-q_i_) ] where f_i_ is the observed frequency for subtype i in the clonotype, and q_i_ is the background frequency of subtype i in the whole sample. To assess statistical significance of measured clonotype bias scores, we generated N random permutations of the observed clonotype data, preserving clonal size and global subtype background frequencies. On the distribution of permuted clonotype bias scores, binned by clone size, we determined expected mean and standard deviation for each clone size bin. Z scores for observed clonotype bias scores were then calculated as the number of standard deviations from the background mean (Z score = 5 corresponds to a p-value ^~^6*10^−7^).

### Statistical analyses for flow cytometry data

Statistical significance was calculated with Prism software. Except where otherwise indicated in figure legends, error bars in graphs indicate standard deviation and statistical comparisons were done by one-way ANOVA test.

### Data availability

Sequence data are deposited in the NCBI Gene Expression Omnibus under accession number GSE182320. Reference atlas can be downloaded (DOI: 10.6084/m9.figshare.16592693) or accessed via the web portal (https://spica.unil.ch/refs/viral-CD4-T). All other data and code are available upon request.

## Acknowledgments

We thank the NIH tetramer facility for reagents; the CCR and University of Rochester Flow Cytometry Core, the University of Rochester Genomics Research Center and the NIH High performance computing cluster for assistance; D. McGavern, T. Mosmann and F. David for technical assistance; R. Bosselut for supporting the research; D. Goldstein, M. Malik and the NCI Office of Science and Technology Resources for their support. We would also like to thank Prof. Carolyn King and David Schreiner at the University of Basel for critical reading of the manuscript. This work was supported by the University of Rochester, and Intramural Research Program of the National Cancer Institute, Center for Cancer Research (CCR), National Institutes of Health, and by the Swiss National Science Foundation (SNF project 180010). The CCR Single Cell Analysis Facility is funded by the Frederick National Laboratory for Cancer Research, Contract HHSN261200800001E. Sequencing was performed with the CCR Genomics Core and the University of Rochester Genomics Research Center.

## Author contributions

M. Andreatta performed research, analyzed data and wrote the manuscript. Z. Sherman and A. Tjitropranoto performed research. M. Kelly contributed to single cell captures and provided guidance. T. Ciucci and S. Carmona supervised the research, analyzed data and wrote the manuscript.

